# Musashi binding elements in Zika and related Flavivirus 3’UTRs: A comparative study *in silico*

**DOI:** 10.1101/407833

**Authors:** Adriano de Bernardi Schneider, Michael T. Wolfinger

**Affiliations:** Department of Medicine, University of California San Diego, 220 Dickinson St, Suite A, San Diego, CA 92103, United States of America; Department of Theoretical Chemistry, University of Vienna, Währingerstraße 17, 1090 Vienna, Austria

## Abstract

Zika virus (ZIKV) belongs to a class of neurotropic viruses that have the ability to cause congenital infection, which can result in microcephaly or fetal demise. Recently, the RNA-binding protein Musashi-1 (Msi1), which mediates the maintenance and self-renewal of stem cells and acts as a translational regulator, has been associated with promoting ZIKV replication, neurotropism, and pathology. Msi1 predominantly binds to single-stranded motifs in the 3’ untranslated region (UTR) of RNA that contain a *UAG* trinucleotide in their core. We systematically analyzed the properties of Musashi binding elements (MBEs) in the 3’UTR of flaviviruses with a thermodynamic model for RNA folding. Our results indicate that MBEs in ZIKV 3’UTRs occur predominantly in unpaired, single-stranded structural context, thus corroborating experimental observations by a biophysical model of RNA structure formation. Statistical analysis and comparison with related viruses show that ZIKV MBEs are maximally accessible among mosquito-borne flaviviruses. Our study addresses the broader question of whether other emerging arboviruses can cause similar neurotropic effects through the same mechanism in the developing fetus by establishing a link between the biophysical properties of viral RNA and teratogenicity. Moreover, our thermodynamic model can explain recent experimental findings and predict the Msi1-related neurotropic potential of other viruses.

## Introduction

Flaviviruses are an emerging group of arboviruses belonging to the *Flaviviridae* family. Researchers have been describing recent outbreaks of these viruses that have not been previously detected for decades^1–3^.

The genus *Flavivirus* comprises more than 70 species that are mainly transmitted by mosquitoes and ticks, typically classified into four groups: Mosquito-borne flaviviruses (MBFVs), tick-borne flaviviruses (TBFVs), insect-specific flaviviruses (ISFVs), that do not have vertebrate hosts, and no known arthropod vector flaviviruses (NKVs), which typically infect bats and rodents. Flaviviruses represent a global health threat, including emerging and re-emerging human pathogens such as Dengue (DENV), Yellow fever (YFV), Japanese encephalitis (JEV), West Nile (WNV), Tick-borne encephalitis (TBEV) and Zika (ZIKV) viruses^4, 5^.

Initially isolated in 1947 from a sentinel rhesus macaque in the Ziika forest, Uganda, ZIKV has not been associated with severe disease, apart from skin rashes, body pain, and fever. Likewise, ZIKV has been circulating across equatorial zones in Africa and Asia for 60 years, until the first outbreak was reported in Yap Island, Micronesia in 2007. Subsequently, the virus spread eastwards to French Polynesia and other Pacific islands in 2013 and reached the Americas in 2015^6, 7^. There are two main ZIKV lineages, the original African (type strain MR766) and an Asian (type strain FSS13025)^8, 9^, the latter also comprising American strains such as PE243.

## Background

The 2015-2017 outbreak in the Americas raised the possibility of a link between ZIKV infection and congenital abnormalities, which included placental damage, intrauterine growth restrictions, eye diseases and microcephaly in children as well as acute motor axonal neuropathy-type Guillain-Barré syndrome in adults ^10^. While MBFVs are typically transmitted by host-vector interaction, vertical transmission from mother to child during pregnancy via transplacental infection has been reported^11^.

The neurotropic potential of ZIKV-related flaviviruses has been known since the 1970s, when Saint Louis encephalitis virus (SLEV) has been attributed to a severe neurological disorder in infected mice^12, 13^. Vertical transmission has been observed with JEV in mice^14^ and human^15^ and a case of human fetal infection have been reported after YFV vaccination^16^. Other transmission pathways of ZIKV include blood transfusions and sexual transmission^17, 18^. Despite enormous efforts in studying ZIKV infections in the last years, the biological reasoning and mechanisms behind arbovirus congenital neurotropism remain elusive.

### Flavivirus genome organization

Flaviviruses have the structure of an enveloped sphere of approximately 50 nm diameter. They are single-stranded positive-sense RNA viruses of 10-12 kb in size, and their genomic RNA (gRNA) encodes a single open reading frame (ORF) flanked by highly structured untranslated regions (UTRs). Upon translation of the ORF, a polyprotein is produced which is processed by viral and cellular enzymes, yielding structured (C, prM, E) and unstructured proteins (NS1, NS2A, NS2B, NS3, NS4A, 2K, NS4B, NS5). Both flavivirus UTRs are crucially related to regulation of the viral life cycle, mediating processes such as genome circularization, viral replication and packaging^19–22^.

### Flaviviruses hijack the host mRNA degradation pathway

The central role of flavivirus 3’UTR in modulating cytopathicity and pathogenicity became apparent when an accumulation of both gRNA and viral long non-coding RNA (lncRNA) has been observed upon infection. These lncRNAs, also known as subgenomic flaviviral RNAs (sfRNAs)^23^, are stable decay intermediates derived from exploiting the host’s mRNA degradation machinery^24^.

sfRNAs are produced by partial degradation of viral gRNA by Xrn1, a host 5’-3’ exoribonuclease that is associated with the endogenous mRNA turnover machinery^25, 26^. The enzyme stalls at highly conserved RNA structures in the viral 3’UTR, so-called Xrn1-resistant RNAs (xrRNAs), resulting in sfRNAs of variable lengths^27, 28^. Xrn1-resistant RNAs and sfRNAs appear to be ubiquitously present in many flaviviruses. They have been described in MBFVs, including DENV^29^, YFV^30^, JEV^31^, and ZIKV^32^, TBFVs^23, 33^, and recently in ISFVs and NKVs^34, 35^. There is typically more than one xrRNA, given the diverse molecular architecture of different flavivirus 3’UTRs. Pseudoknot interactions have been proposed in some, but not all flavivirus xrRNAs^32, 36^. While they may form transiently under certain conditions^28^, conclusive validation of their ubiquitous presence is missing. Hence, we will exclude them in this work. Earlier studies in our group have identified conserved RNA structural elements in viral 3’UTRs ^37–41^, some of which have later been attributed to xrRNA functionality^23^. Stem-loop (SL) as well as dumbbell (DB) structures are found in 3’UTRs of flaviviruses in single or double copies (Figure 1) and have been associated with quantitative protection of downstream viral RNA^42^.

**Figure 1.**
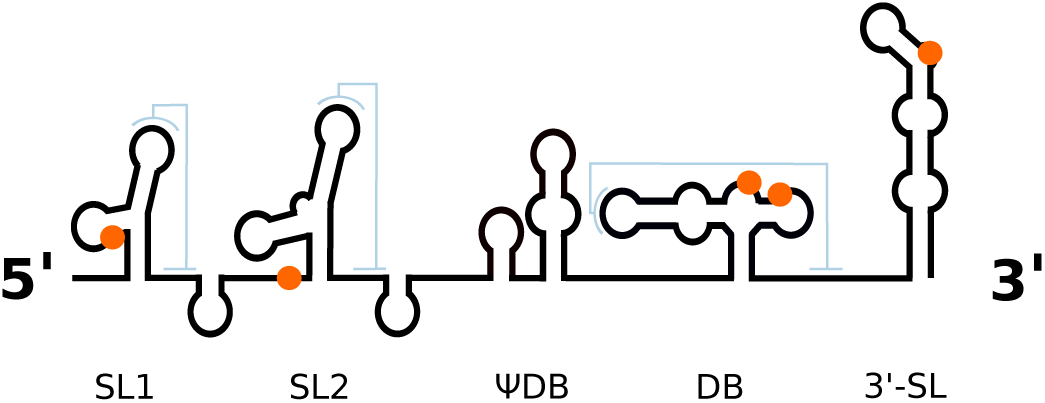
Schematic representation of the ZIKV 3’UTR. Conserved RNA elements include two stem-loop structures (SL1 and SL2), a ΨDB and canonical DB element as well as the terminal 3’ stem-loop structure (3-SL). Positions of Musashi-binding *UAG* motifs in the Asian/American ZIKV lineage are highlighted in orange. Possible pseudoknot interaction sites (sketched in light blue) do not overlap with potential Musashi binding sites.

The inhibition of Xrn1 by viral RNA yields sfRNAs that affect many cellular processes, both in the vector and the host^43^. In mosquitoes, sfRNA interacts directly with the predominant innate immune response pathway, RNA interference (RNAi), by serving as a template for microRNA (miRNA) biogenesis^44^. Conversely, in host cells sfRNA modulates the anti-viral interferon response^45^, e.g., by binding proteins to inhibit the translation of interferon-stimulated genes^46^. Moreover, sfRNA has been shown to inhibit Xrn1 and Dicer activity, thereby altering host mRNA levels^47, 48^.

At the same time, a variety of host proteins bind the 3’UTR of flaviviruses, thereby mediating viral replication, polyprotein translation or the anti-viral immune response (see Table 1 in ref.^43^ for a comprehensive overview of host proteins that bind flavivirus 3’UTR/sfRNA). Although notoriously underrepresented in literature, one can expect that many of these proteins also bind sfRNA due to sequence and structure conservation.

**Table 1.**
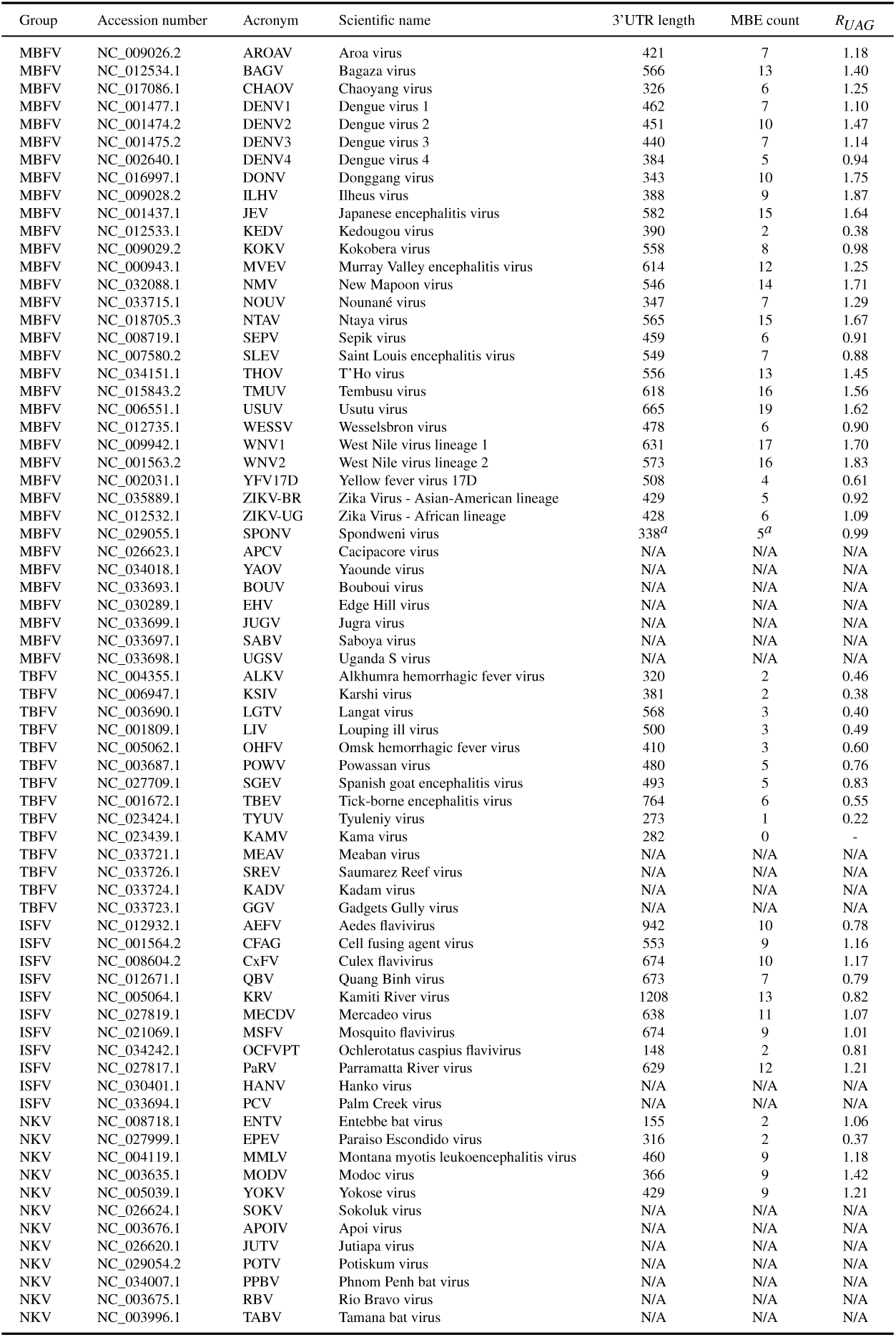
Viral genomes analyzed in this study. Flaviviruses are categorized into the groups mosquito-borne flaviviruses (MBFV), tick-borne flaviviruses (TBFV), insect-specific flaviviruses (ISFV) and no known vector flaviviruses (NKV). 3’UTR lengths, number of Musashi binding elements found in 3’UTRs and relative MBE abundance *R*_*UAG*_ are listed. *R*_*UAG*_ values above 1 indicate relative enrichment of *UAG* trinucleotides, whereas values below 1 indicate relative depletion. ^*a*^3’UTR length and MBE count of SPONV SA-Ar strain. ^*N/A*^3’UTR partial or not available in the `refseq` data set.

### Subgenomic flaviviral RNA interacts with Musashi

One of these groups of host factors is the Musashi (Msi) protein family. Msi is a highly conserved family of proteins in vertebrates and invertebrates that act as a translational regulator of target mRNAs and is involved in cell proliferation and differentiation. While the two Msi paralogs in mammals, Musashi-1 (Msi1) and Musashi-2 (Msi2), are expressed in stem cells^49–51^ and overexpressed in tumors and leukemias^52^, they are absent in differentiated tissue. Moreover, Msi1 is involved in the regulation of blood-testis barrier proteins and spermatogenesis in mice^53^. Musashi proteins have two RNA recognition motif (RRM) domains, whose sequence specificity has been determined by an *in vitro* selection method and NMR spectroscopy^51, 54, 55^. The trinucleotide sequence *UAG*, whose thermodynamic binding specificity was determined by fluorescence polarization assays, has been identified as core Musashi binding element (MBE). Nucleotides enclosing the main MBE recognition motif make minor contributions to binding affinity^56^. While earlier SELEX experiments identified the binding aptamer sequence (*G/A*)*U*_*n*_*AGU* (*n* = 1 – 3)^51^, iCLIP experiments with Msi1 in human glioblastoma cells confirmed the preferential binding of Msi1 to single-stranded (stem-loop) *UAG* sequences in 3’UTRs, but not in coding regions^57^. Zearfoss et al.^56^ observed that both *GUAGU* and *AUAGU* are recognized by mouse Msi1, whereas Drosophila Msi1 has a higher affinity for *GUAGU*. NMR-derived structures of the two Msi1 RNA recognition motifs in complex with RNA also show that both RNA-binding domains bind *GUAGU* (PDB IDs 2RS2 and 5X3Z).

In summary, there is a strong consensus in the literature that *UAG* is central to all proposed Musashi binding motifs. Therefore, we focus our calculations around this trinucleotide, and provide evidence that the availability of *UAG* in pentanucleotides expands to the accessibility of the entire motif.

### Musashi is involved in Flavivirus neurotropism

An interesting, yet understudied hypothesis is the possibility that the stem cell regulator protein Musashi could be related to ZIKV tropism. Based on the identification of a MBE in the 3’UTR of the ZIKV genome^10^, de Bernardi Schneider et al.^7^ reported the presence of the same element with a higher binding affinity for human Msi1 in all ZIKV sequences that belong to the Asia-Pacific-Americas clade in an *in silico* screen and implied that there could be a change of tropism for the viral lineage. Chavali et al.^58^ tested the possibility of Msi1 interaction with the ZIKV genome *in vivo* and found that Msi1 not only interacts with ZIKV, but also enhances viral replication. They noted that ZIKV RNA could compete with endogenous targets for binding Msi1 in the brain of the developing fetus, thereby dysregulating the expression of genes required for neural stem cell development. Based on their data the authors concluded that Msi1 is involved in ZIKV neurotropism and pathology and raised the question whether MBEs present in other flavivirus genomes could exhibit similar functionality. In a recent study, Platt et at.^59^ investigated whether ZIKV-related arboviruses can cause congenital infection and fetal pathology in utero in immunocompetent mice. They tested two emerging neurotropic flaviviruses, WNV, and Powassan virus (POWV), as well as two alphaviruses, Chikungunya virus (CHIKV) and Mayaro virus (MAYV). All four viruses caused placental infection, however, only WNV and POWV resulted in fetal demise, indicating that ZIKV is not unique among flaviviruses in its capacity to be transplacentally transmitted and cause fetal neuropathology.

In this contribution, we systematically analyze the Musashi-related neurotropic potential of well-curated flavivirus genomes *in silico*. We investigate structural features of MBEs in viral 3’UTRs by a thermodynamic model of RNA structure formation and work out the biophysical properties of conserved RNA structures harboring MBEs in order to build a theoretical ground for future *in vivo* studies.

## Materials and Methods

### Dataset

Sequence data for the present study was acquired from the public National Center for Biotechnology Information `(NCBI) refseq` database (https://www.ncbi.nlm.nih.gov/refseq/) on 15 December 2017. We filtered for all complete viral genomes under taxonomy ID 11051 (genus *Flavivirus*), resulting in 72 genomes, 51 of which had 3’UTR sequences and annotation available (Table 1).

The core Musashi binding element is only three nucleotides long, hence one can expect to observe a certain number of *UAG* trinucleotides by chance in any viral 3’UTR. Table 1 shows the number of MBEs present in 3’UTR regions of viral genomes analyzed here as well as the ratio *R*_*UAG*_ = *O*_*UAG*_/*E*_*UAG*_, i.e., observed versus expected frequencies. Assuming that all four nucleotides (*A,U,G,C*) occur independently and with equal probability, the expected probability to observe a subsequence of length *l* is equal to (1/4)^*l*^. For *l* = 3, this is equal to 1/64. More realistically, the frequency of each nucleotide *i* ∈ {*A,U,G,C*} in an RNA sequence of length *L* is *F*_*i*_ = *N*_*i*_/*L*, where, *N*_*i*_ is the nucleotide count of *i*. For any trinucleotide *XYZ*, the expected trinucleotide frequency *E_XY Z_* is then computed from mononucleotide frequencies as *E*_*XYZ*_ = *F*_*X*_ ∗ *F*_*Y*_ ∗ *F*_*Z*_.

The `refseq` genome for Spondweni virus (SPONV, accession number NC_029055.1) does not include a 3’UTR sequence. Since SPONV is phylogenetically closely related to ZIKV^60^, we were looking to include this sequence into our analysis. Nikos Vasilakis (Univ. of Texas Medical Branch, Galveston, TX, USA) generously provided SPONV sequence data. The 338 nt 3’UTR sequence of the SA-Ar strain (see supplementary material) has been added to the set of flavivirus sequences analyzed here.

Kama virus (KAMV) does not contain *UAG* trinucleotides in the 3’UTRs, consequently it has been discarded from our dataset. The remaining virus species contain between 1 and 19 MBEs in their 3’UTRs.

### Opening energy directly relates to single-strandedness

The biophysical model employed here is based on a description of RNA at the level of secondary structures, building upon the thermodynamic *nearest neighbor* energy model as implemented in the `ViennaRNA` Package^61^. This allows for computing equilibrium properties of RNA such as the single most stable, minimum free energy (MFE) structure, as well as the partition function 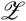. The latter makes an evaluation of the thermodynamic ensemble of RNA structures available and is defined as the sum over all Boltzmann factors of individual structures *s*

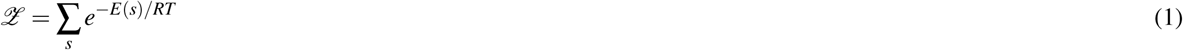

where *E*(*s*) is the free energy of the structure, *R* the universal gas constant and *T* the thermodynamic temperature of the system. The equilibrium probability of a secondary structure *s* is then defined as

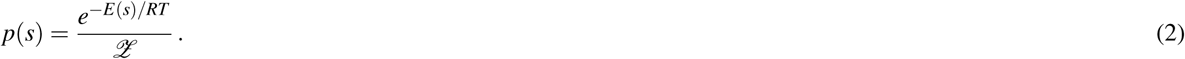

The partition function 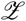 can be computed efficiently via dynamic programming^62^ and allows calculation of individual base pair probabilities, even for large sequences^63^. In this line, the *accessibility* (i.e., the probability that a region *i…j* along the RNA is single-stranded) can be derived from the partition function (Eq. 1)^64^. Likewise, the *opening energy* (i.e., the free energy required to force the region to be single-stranded) can be computed as

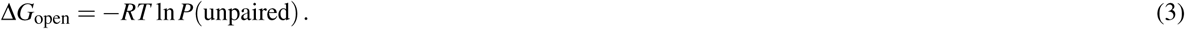

The opening energy of a region within an RNA is directly related to local RNA secondary structure. In this line, low opening energy is a reliable indicator for single-strandedness. We employ the sliding window approach of `RNAplfold`^63^ to compute local pairing probabilities of *UAG* trinucleotide motifs to assess the likelihood of single-strandedness of and around MBEs. `RNAplfold` is part of the `ViennaRNA` Package^61^ and can compute the accessibilities or single-strandedness of all intervals of an RNA in cubic time^65^. We select 97 nt windows upstream and downstream of MBEs in viral 3’UTRs and compute local pairing probabilities for base pairs within 100 nt windows. Opening energies for trinucleotides are then evaluated from averaged pairing probabilities with `RNAplfold`.

The significance of a calculated MBE opening energy is assessed by comparison with a large number of randomized sequences of the same length and same base or dinucleotide composition. We compute the opening energies of trinucleotides both in a genomic as well as a shuffled sequence context and apply a *z* score statistics. The normalized *z* score is defined as

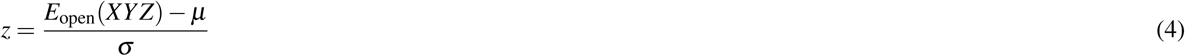

where *E*_open_(*XYZ*) is the opening energy of trinucleotide *XYZ* in its genomic context, *µ* and *σ* are the mean and standard deviations, respectively, of the opening energies of *XYZ* computed over a large sample of randomized sequences. Randomization with regard to keeping sequence composition is achieved here by applying dinucleotide shuffling to the 97 nt windows upstream and downstream of MBEs, while keeping *XYZ* in place. The same idea applies to calculations of pentanucleotide motifs.

The approach outlined above is implemented in the Perl utility `plfoldz.pl`, which is available from https://github.com/mtw/plfoldz. The script employs the `ViennaRNA`^61^ scripting language interface for thermodynamics calculations, the `ViennaNGS`^66^ suite for extraction of genomic loci and the uShuffle Perl bindings^67^ for k-let shuffling. The tool reports for each requested trinucleotide the opening energy in a genomic context as well as an opening energy *z* score obtained from *n* shuffling events of upstream and downstream sequences. Here, *n* = 10,000 dinucleotide shuffling events were used.

### Characteriztaion of MBEs within xrRNAs

To localize MBEs within homologous substructures in flavivirus 3’UTRs we constructed `infernal`^68^ covariance models for conserved xrRNA elements. The structural RNA alignments underlying the `infernal` models were computed with `locarna`^69^ and further analyzed with `RNAalifold`^61^ and `RNAaliSplit`^70^.

## Results

### MBEs are highly accessible in ZIKV 3’UTRs

The Musashi family of proteins preferentially bind single-stranded *UAG* motifs in 3’UTRs^57^. To evaluate the thermodynamics of Msi-*UAG* affinity more broadly, we set out to analyze the single-strandedness of all possible trinucleotides in ZIKV genomes. To this end, we computed the opening energies of all trinucleotide motifs present in the coding sequence (CDS) and 3’UTR of the African (ZIKV-UG) and Asian/American (ZIKV-BR) Zika strains. A *z* score was calculated for each occurrence of trinucleotide *XYZ* according to Eq. 4, thereby normalizing the opening energy of *XYZ* in its genomic context with *n* = 10,000 dinucleotide-shuffled upstream and downstream regions of 97 nt, using 100 nt windows in `RNAplfold`.

Negative opening energy *z* scores indicate increased accessibility, i.e., *UAG* trinucleotides in viruses with overall low *z* scores are likely to occur in an unpaired structural context within the 3’UTR. Through the distribution of *z* scores, sorted by median *z* score (Figure 2) we were able to see three aspects standing out. First, the distribution of *z* scores is markedly divergent among CDS and 3’UTR. The interquartile ranges of opening energy *z* scores are homogeneous within the CDS region, while dispersion is varied within the 3’UTR. We hypothesize that this is caused by a different sequence composition that manifests in highly variable opening energies. It could, however, also be an artifact of the different sample sizes based on the divergent trinucleotide count in CDS and 3’UTR, respectively. Second, *UAG* is the most accessible trinucleotide in the 3’UTR of ZIKV-BR and among the highest accessible trinucleotides in the 3’UTR of ZIKV-UG. This is striking as it corroborates previous experimental evidence of Musashi affinity to ZIKV^58^ by means of a thermodynamic model, thus underlining a possible role of Msi1 in ZIKV neurotropism. Moreover, the *UAG* trinucleotide is neither enriched nor depleted in the 3’UTRs of ZIKV-BR and ZIKV-UG (Table 1). Third, the canonical start codon *AUG* appears to the far right end of the scale in both ZIKB-BR and ZIKV-UG 3’UTRs, i.e., it is among the least accessible trinucleotides. This suggests evolutionary pressure on keeping the start codon in a paired structural context within the 3’UTR, thereby prohibiting accessibility to ribosomes and disabling undesirable leaky translation start from these *AUG* triplets.

**Figure 2.**
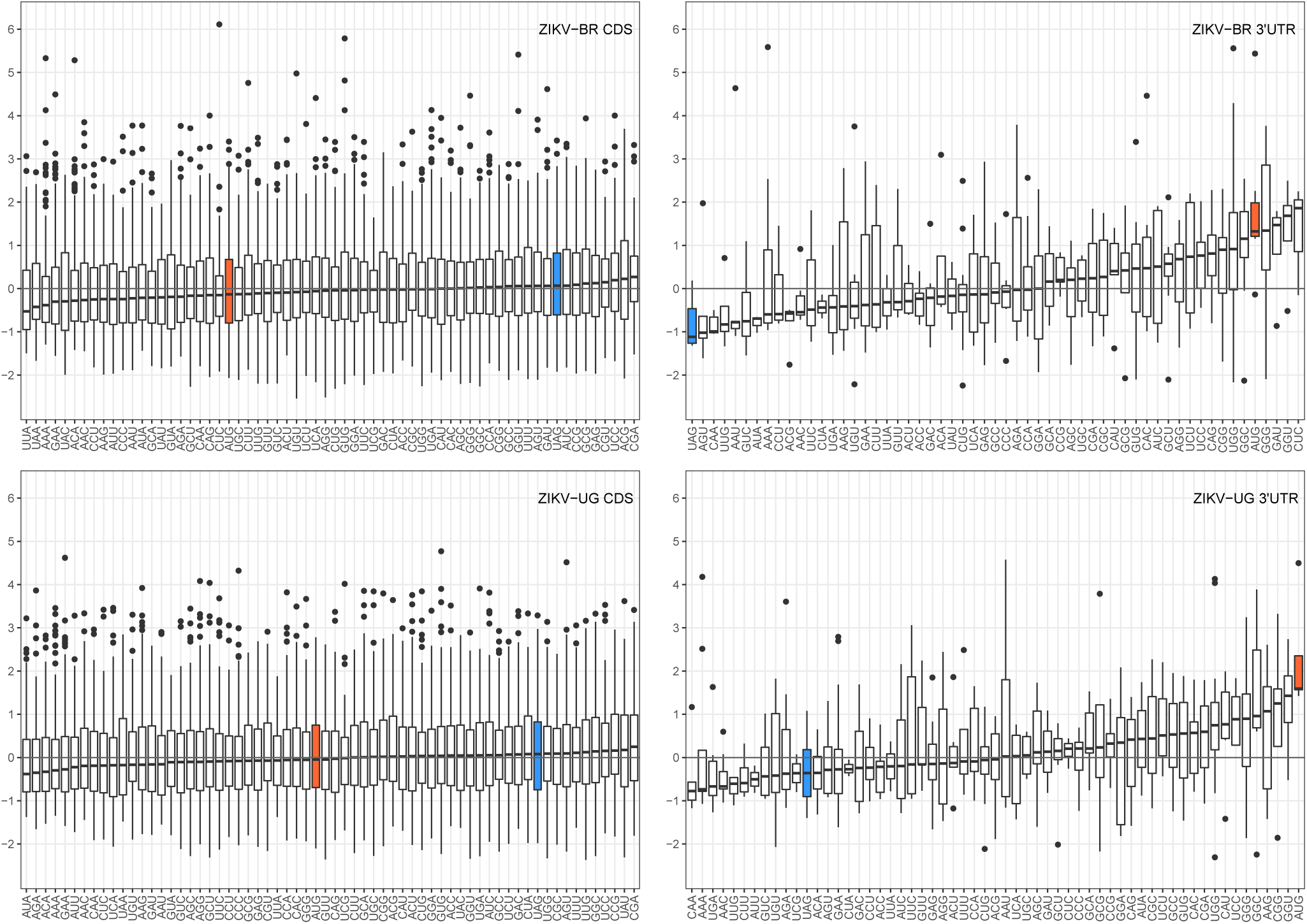
Distribution of *z* scores of opening energies for trinucleotides found in the coding region (CDS, left) and 3’UTR (right) of ZIKV from Brazil (top) and Uganda (bottom), sorted by median opening energy *z* score. The MBE motif is highlighted in blue and shows low overall *z* scores in the 3’UTR, indicating that this trinucleotide is more likely to appear in a single-stranded structural context. Contrary, the canonical start codon *AUG* (highlighted in orange) shows high opening energy *z* scores, indicating reduced accessibility within the 3’UTR region. Data for trinucleotides *AUU*, *UAA*, *UCG* and *UUU* are omitted because they only occur once within the 3’UTR of ZIKV-BR. Likewise, trinucleotides *CGU*, *GUA*, *UAA* and *UAU* are omitted in the ZIKV-UG 3’UTR plot.

We also tested the accessibility of larger Musashi recognition motifs. To this end, we employed the same approach outlined above for all pentanucleotides found in the 3’UTRs of ZIKV-BR and ZIKV-UG, respectively. The distributions of opening energy *z* scores (supplementary data, Figure S1) are in good agreement with our results derived for trinucleotides as well as previous experimental data, suggesting that core a *UAG* is among the most accessible motifs. In particular, our data shows that *NUAGN* is the most accessible pentanucleotide in the 3’UTR of ZIKV-BR and among the highest accessible pentanucleotides in the 3’UTR of ZIKV-UG, similar to the situation found for trinucleotides. *UAG* appears to be conserved in an unpaired structural context not only by itself but also in a larger sequence context of enclosing nucleotides, which exhibit high accessibility upon the presence of a central *UAG* Musashi recognition element. This finding is in line with the reported Musashi recognition pentamers *GUAGU* and *AUAGU*.

### MBE accessibility in related viruses

To assess the Musashi-related neurotropic potential of other flaviviruses, we evaluated the accessibilities of MBEs in related species. To this end, all 435 *UAG* trinucleotide motifs within 3’UTRs in the `refseq` dataset were identified, grouped by vector specificity and subjected to the computational approach outlined above (97 nt upstream/downstream windows, *n* = 10,000 dinucleotide shufflings).

Msi1 preferentially binds single stranded RNA^57^, consequently *UAG* motifs that contribute with low *z* scores have a high affinity for Msi1 binding. Within the MBFV group, the Asian/American lineage Zika virus (ZIKV-BR) has the lowest median *z* score, followed by Saint Louis encephalitis virus (SLEV), Nounané virus (NOUV) and the African lineage Zika virus (ZIKV-UG). Among others, two lineages of West Nile virus (WNV1, WNV2) and Yellow fever virus (YFV) appear with a negative median *z* score. ZIKV-BR turns out to be the only isolate among MBFVs that has just negative *z* score values in our simulations, i.e., all *UAG* motifs within the 3’UTR of the Brazilian ZIKV isolate appear in an unpaired structural context. Likewise, Karshi virus (KSIV), Alkhumra hemorrhagic fever virus (ALKV) and Langat virus (LGTV) have a strictly negative *z* scores distribution among the TBFV group. POWV, Omsk hemorrhagic fever (OHFV) and Louping ill virus (LIV) have negative mean opening energy *z* scores. Interestingly, *UAG* trinucleotides are relatively depleted in all TBFV species analyzed here (Table 1). Among NKVs, Montana myotis leukoencephalitis virus (MMLV) and Entebbe bat virus (ENTV) show negative mean opening energy *z* scores. Culex flavivirus (CxFV), Cell fusing agent virus (CFAG), Parramatta River virus (PaRV) and Ochlerotatus caspius flavivirus (OCFVPT) tend to have singe-stranded MBEs among the ISFVs. Here, OCFVPT is the only isolate with a strictly negative *z* score distribution (Figure 3).

**Figure 3.**
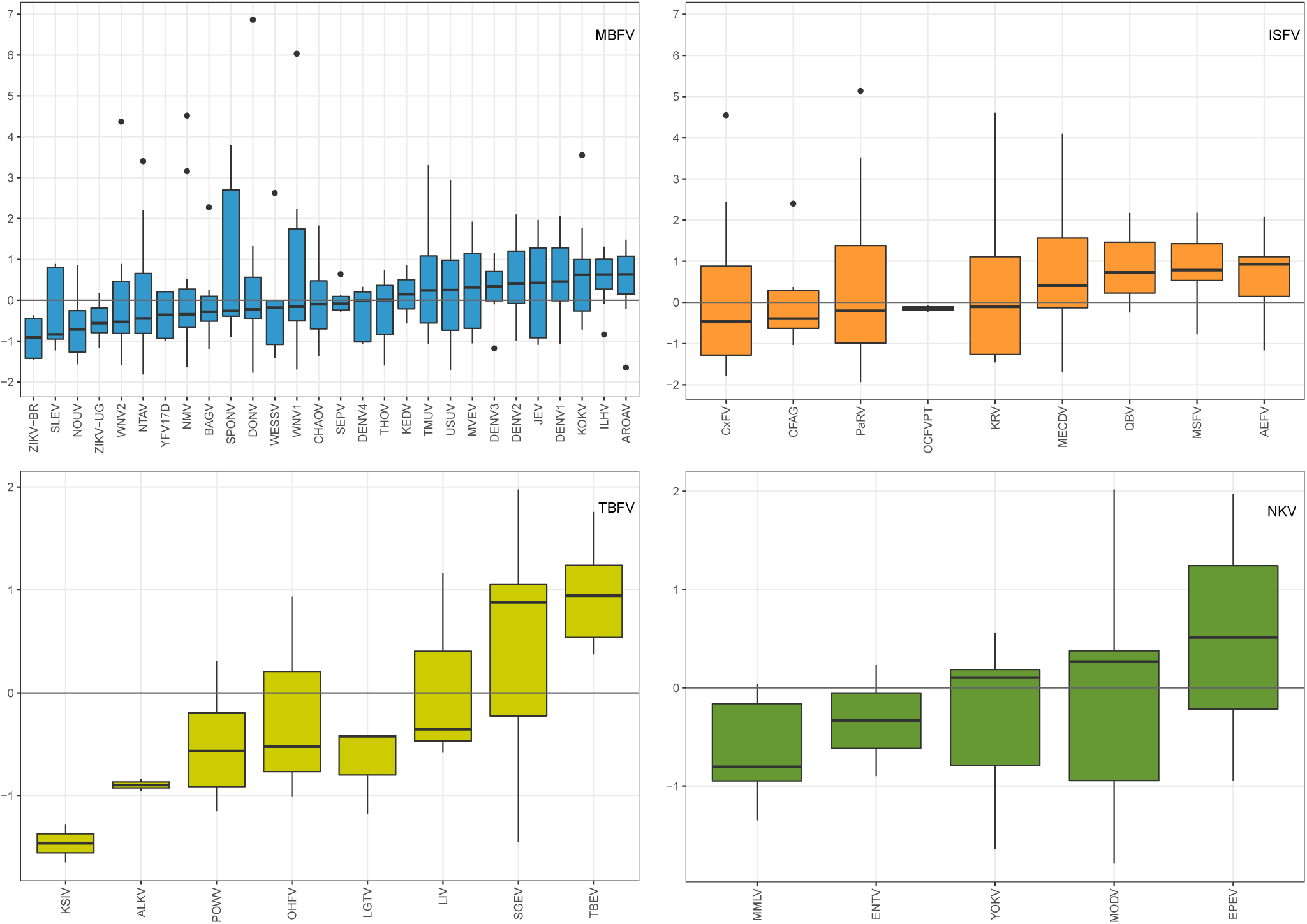
Distribution of MBE opening energy *z* scores in flavivirus 3’UTRs, grouped by vector specificity and sorted by median *z* scores. Top left: MBFVs, top right: ISFVs, bottom left: TBFVs and bottom right: NKVs. The Asian/American lineage ZIKV-BR isolate has the lowest median *z* score among all MBFV. Alkhurma virus (ALKV), Ochlerotatus caspius flavivirus (OcFV), ENTV and EPEV contain only two MBEs on the 3’UTR. Tyuleniy virus (TYUV) was excluded as it only contains a single MBE.

The number of *UAG* trinucleotide motifs in 3’UTRs of the `refseq` dataset lies between 1 and 19 (Table 1). The overall range of opening energy *z* scores is not equal among different flavivirus groups. While the lower bound is between –1.65 and –1.93 among all groups, MBFVs and ISFVs show markedly higher upper bounds than TBFVs and NKVs, respectively. Absolute values of computed MBE opening energy *z* scores are listed in Table 2.

**Table 2.** Distribution ranges of opening energy *z* scores for MBEs in the 3’UTR of flaviviruses. The minimal and maximal *z* score is listed for all *UAG* motifs (total) within a the 3’UTR, and only for those that overlap one of the conserved xrRNA elements SL and DB. Dashes indicate that SL/DB elements are not conserved.

|  | total |  | SL |  | DB |  |
| --- | --- | --- | --- | --- | --- | --- |
|  | min | max | min | max | min | max |
| MBFV | -1.82 | 6.86 | -1.35 | 2.10 | -1.42 | 2.23 |
| ISFV | -1.93 | 5.14 | - | - | - | - |
| TBFV | -1.65 | 1.97 | - | - | - | - |
| NKV | -1.79 | 2.02 | - | - | -1.79 | 2.02 |

### Conserved xrRNAs contain MBEs

Several species appear to the right of the plots in Figure 3 due to the sorting by median *z* score. However, they comprise a non-negligible number of accessible MBEs, as indicated by negative opening energy *z* scores. Examples are (re-) emerging species like JEV and Usutu virus (USUV), which contain 15 and 19 MBEs, respectively.

To investigate this further, we assigned each MBE in our dataset to one of the conserved elements stem-loop (SL), dumbbell (DB) and 3’ stem-loop (3SL) (Figure 1) by means of covariance models. Analysis of RNA sequence and structure conservation revealed that the majority of virus isolates in our dataset contain only a single *UAG* motif within their SL and 3SL elements. Conversely, DB elements, which are conserved in MBFVs and NKVs (Figure 4), stand out among conserved RNA structures in flavivirus 3’UTRs. They contain a pair of MBEs, separated by a 4 nt spacer, within a perfectly conserved sequence motif of approx. 20 nt length in their distal stem-loop structure. We hypothesize that this pair of conserved *UAG* motifs interacts with the two RNA-binding domains in Musashi proteins. Figure 5 shows the consensus secondary structure of flavivirus DB elements.

**Figure 4.**
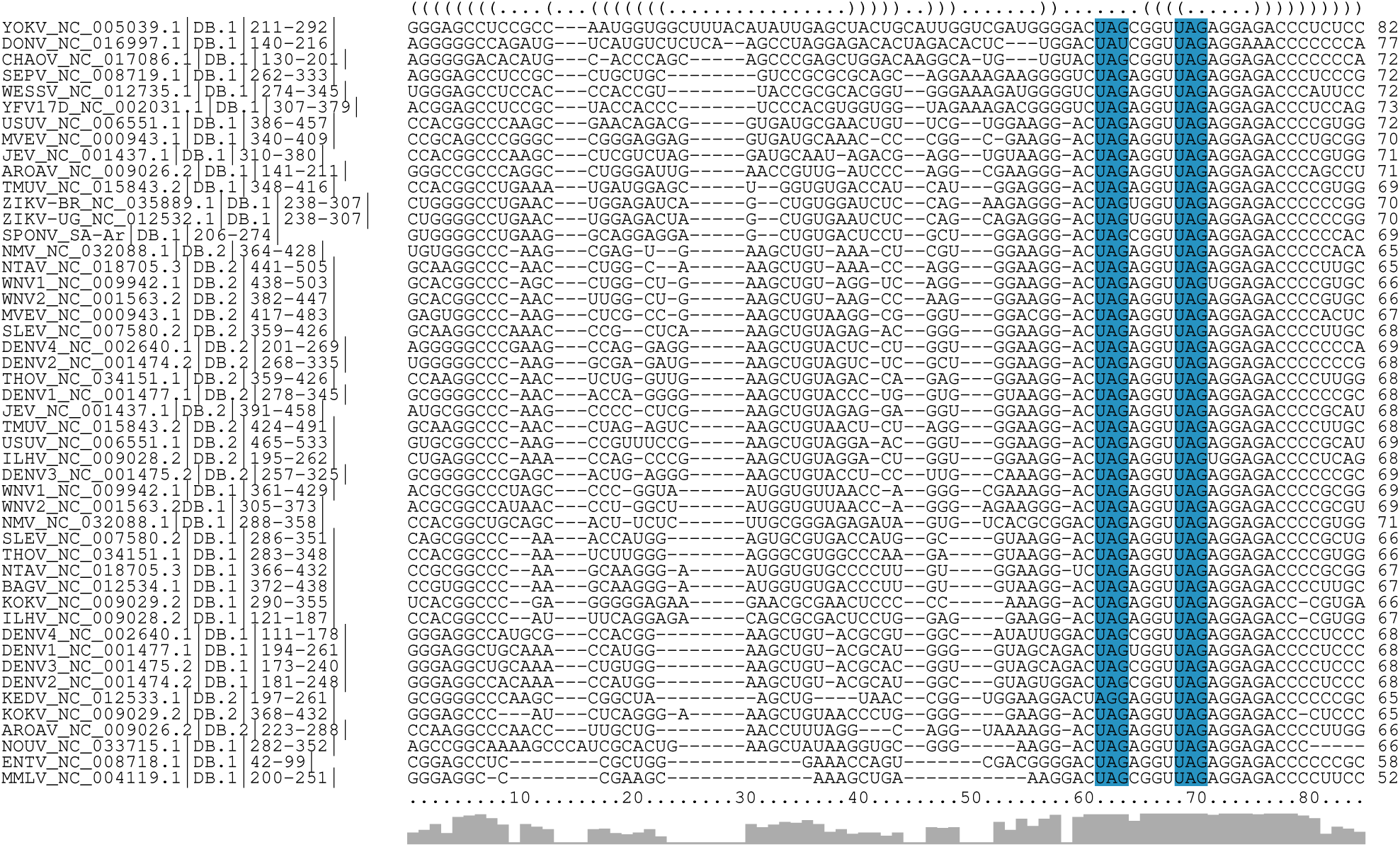
Structural alignment of conserved dumbbell (DB) elements in the 3’UTR of MBFV and NKV flaviviruses. Several species have two copies of DB elements in their 3’UTR, indicated by `DB.1` and `DB.2` in the sequence identifier. Coordinates are given relative to the 3’UTR start. A consensus structure in dot-bracket notation is plotted on top of the alignment. Gray bars at the bottom indicate almost perfect sequence conservation within the distal stem-loop sub-structure (positions 60-80). Two conserved MBE motifs in the central multiloop and distal stem loop are highlighted in blue.

**Figure 5.**
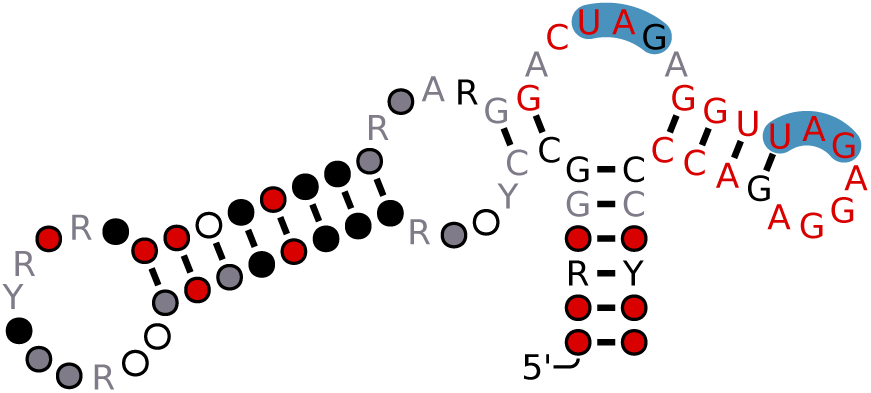
Consensus secondary structure of the flavivirus DB element with MBEs highlighted in blue. Figure generated from the MSA in Figure 4 with `R2R`^71^. Structure and color annotation inferred by `R2R`. Nucleotide symbols represent conserved nucleotides. Circles represent columns in the MSA that are typically or always present but do not conserve nucleotide identity. Red, black and gray colors indicate the level of nucleotide conservation, in decreasing order.

## Discussion

Our findings lead to the conclusion that the accessibility of *UAG* motifs calculated through opening energies in Flavivirus 3’UTRs is indicative of the Musashi-related neurotropic potential of virus species. Our computational analyses show that there is little difference in the distribution of opening energies for all trinucleotides within the polyprotein (CDS) region of ZIKV. When comparing CDS and 3’UTR regions, we see a difference in behavior, as different trinucleotides do possess different opening energies, *UAG* being highly accessible in ZIKV. Although not possible to quantify the impact of the accessibility on the patient phenotype, it is interesting to see the *UAG* motifs on the Brazilian ZIKV isolate more accessible than on the Ugandan ZIKV isolate. This result raises the question once again if the increased pathogenicity seen in ZIKV today is due to changes in the sequence over time or simply lack of better surveillance^72^.

Previous experiments lead toward the idea that ZIKV is unique among flaviviruses regarding the clinical outcomes resulting from congenital infection^73, 74^. Although our results indicate that this may be true for well-studied viruses such as DENV and WNV, other viruses which have not caused recent outbreaks may have been neglected.

Looking in depth at other viruses, Nounané virus (NOUV), a dual-host affiliated insect-specific flavivirus is found among the viruses with high MBE accessibility. NOUV was isolated in Cote d’Ivoire in 2004 from *Uranotaenia mashonaensis*, a Culicidae mosquito not known to harbor flaviviruses before^75^. While replication has been tested on human and non-human cell lines, vertebrate infection and pathogenesis could not be observed^76^.

Within the TBFV serocomplex, KSIV has the lowest overall MBE opening energies. Originally isolated from *Ornithodoros papillipes* ticks in Uzbekistan in 1972^77^ it currently does not present history of infection in humans. Conversely, Powassan virus (POWV), another TBFV with negative MBE opening energy, first isolated in Powassan, Ontario, Canada in 1959 from a child who died of acute encephalitis^78^ can cause transplacental infection^59^ and has been associated with severe neuropathology and death in mice and human^79^.

Given that *UAG* can be regarded as the primary Msi1 binding motif, we can argue that ZIKV has the highest affinity for binding Msi1 among all MBFVs. Platt et al.^59^ showed that besides ZIKV, the neurotropic flaviviruses WNV and POWV, as well as the alphaviruses Chikungunya (CHIKV) and Mayaro (MAYV) infect placenta and fetus in immunocompetent, wild-type mice. However, only WNV was shown to infect the placenta and the fetal central nervous system, causing injury to the developing brain.

Congenital infection in humans is documented for WNV, JEV, YFV, and ZIKV. CHIKV and MAYV did not show this behavior. In this line, our results are in agreement with experimental studies that reported teratogenicity for SLEV ^12,13^, WNV^59,80^, YFV^16,81^ and POWV^59^.

Bizarre neurological manifestations were also observed in patients infected by Ntaya virus (NTAV)^82^, a neurotropic virus from the Japanese encephalitis serocomplex, as well as WNV in humans ^83,84^ and in mice^85^, USUV ^86,87^ and DENV ^88,89^. The fact that these viruses line up more on the positive side of the opening energy plots in Fig. 3 does not mean that they should not be neurotropic. It merely highlights that there might be additional mechanisms causing neuropathogenicity.

### MBEs are conserved in flavivirus 3’UTR elements

Flavivirus DB elements do not only show structural conservation over the MBFV and NKV serocomplexes, but even maintain their primary sequence within a region of approx. 20 nt of the distal stem-loop (Figs. 4 and 5). The combination of covariation and primary sequence conservation within a single RNA element underlines the importance of DB elements in flavivirus pathogenicity. It could also be indicative of a special role of DB element regions in the minus-strand synthesis during flavivirus replication.

*UAG* trinucleotides are the core nucleotides within MBE motifs to contribute the highest binding energy^56^. Our current analysis underlines that there seems to be evolutionary pressure on keeping *UAG* motifs within the DB elements unpaired. In ZIKV we see that not only the *UAG*s within DB elements but also those that overlap with SL elements show negative opening energy *z*-score.

### Msi1 presence in different cells

The presence of Msi1 proteins in both sperm and neural precursor cells highlights the importance of studying the Msi1-MBE interaction in flaviviruses. Given that Msi1 has been shown to enhance ZIKV replication^58^, this interaction could be a critical reason why ZIKV persists in sperm for a long time after the individual has been infected^90,91^, allowing the virus to be transmitted sexually and also why the virus would harbor itself in neuronal cells, allowing it to interfere with and dysregulate neurodevelopment.

### A possible role of Musashi in the flavivirus life cycle

Msi1, which binds to the 3’UTR of target mRNAs, has been shown to repress translation initiation by competing with the translation initiation factor eIF4G for binding to poly(A)-binding protein (PABP), thereby inhibiting the assembly of the 80S ribosomal unit^92^. Ribosome profiling experiments have corroborated this down-regulatory effect of Msi1, while keeping mRNA levels^93^. This allows for a speculative explanation of the findings by Chavali et al.^58^, i.e., that Msi1 enhances ZIKV replication, and a possible role of Msi1 in the viral life cycle: Flaviviruses need to “donate” a few copies of the quasispecies ensemble for Xrn1 degradation and subsequent sfRNA production. In this line, Msi1 could serve as an agent that provides a reasonable amount of gRNAs that are not translated but subject to Xrn1 degradation. The resulting sfRNAs can then down-regulate the host response^46,94^.

## Conclusion

We studied a specific aspect of flavivirus congenital pathogenicity, i.e., the neurotropic effect inferred by the presence of MBEs in the 3’UTR of flavivirus genomes. Employing an established biophysical model of RNA structure formation, we analyzed the thermodynamic properties of MBEs *in silico*. Our results underline experimental studies suggesting that ZIKV is not alone in its capacity to cause severe neuropathology to infants through the MBE mechanism. While several tick-borne and mosquito-borne flavivirus species like Karshi virus (KSIV), Alkhumra hemorrhagic fever virus (ALKV) or Nounané virus (NOUV) line up with ZIKV in our theoretical model, their tropism might have been overseen due to the lack of reported significant outbreaks. However, some of them appear to have similar neurotropic potential and thus might be potent emerging pathogens.

The approach presented here could in principle be used for developing a tool to predict the Musashi-related neurotropic potential of novel viruses or (re-)emerging strains of known viruses. Combination of opening energy *z* scores with large scale epidemiologic data could be employed in a machine learning framework that also considers structural conservation and homology of flavivirus 3’UTR elements. Such a tool could play a role in categorizing viruses.

## Acknowledgements

We thank Nikos Vasilakis for providing Spondweni virus sequences. We further thank Ivo Hofacker for fruitful discussions.

## Author contributions statement

A.B.S. and M.T.W. conceived the study, conducted the *in silico* experiments, analysed the results and wrote the manuscript. Both authors reviewed the manuscript.

## Data availability

The `plfoldz.pl` Perl Utility for computing RNA opening energy *z* scores is available from https://github.com/mtw/plfoldz.

