## Supplementary data for "Musashi binding elements in Zika and related Flavivirus 3’UTRs: A comparative study *in silico*"

### 1 Pentanucleotide accessibility in ZIKV 3'UTR

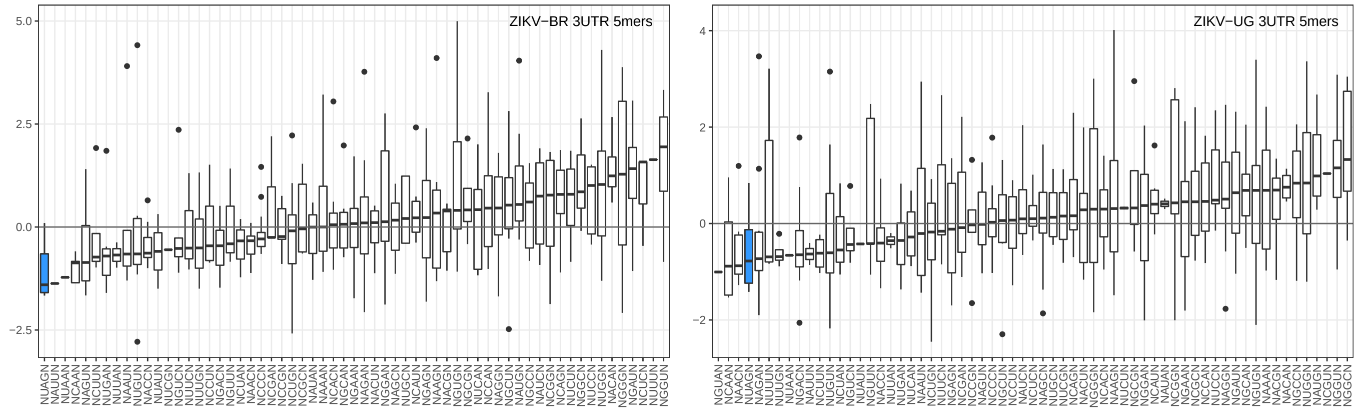


Figure 1: Distribution of $z$ scores of opening energies for pentanucleotides found in the 3'UTR of ZIKV from Brazil (left) and Uganda (right), sorted by median opening energy $z$ score. Each bin comprises data for all pentanucleotides $NXYZN$ that have a central trincleotide $XYZ$ enclose by arbitrary nucleotides $N$. Pentanucleotides $NUAGN$ (highlighted in blue) have low opening energy $z$ scores, i.e. they are highly accessible in the structural ensembles of both ZIKV lineages.

### 2 Spondweni virus 3'UTR

The downloaded `refseq` genome for Spondweni virus (SPONV) `NC_029055.1` does not include the 3'UTR sequence. Since SPONV is phylogenetically related to Zika virus (ZIKV) [1], we were looking to include this sequence into our analaysis.

Nikos Vasilakis (Univ. of Texas Medical Branch, Galveston, TX) generously provided SPONV sequences. The 338 nt 3'UTR sequence of the SA-Ar strain (listed below) has been added to the set of flavivirus sequences analyzed in this study.

```
>SPONV SA-AR|3UTR|
AUAAUGUAAAUAAUAAAUAAAUUAAUAAAGUAAGGAUAGGAAACUAACCUAGCCUAACUA
ACAAAGUCAGGCCGUAGAUUAAGACGCCAUGGCACGGAAGAAGCCAUGCUGCCUGUGAGC
CCCCCAGGAGGAUCUGGGUUAACAAAGAGAGCAAUGUCUCUCCCACGCCUGGAAGAGGUG
GCGAUCUCUCCAGAGCGGUAAAAGCGUGGGGCCUGAAGGCAGGAGGAGCUGUGACUCCUG
CUGGAGGGACUAGCGGUUAGAGGAGACCCCCCACAAAACGCAAAACAGCAUAUUGACGCU
GGGAAAGACCAGAGACUCCGUGCGUUUCCAGCACGCCG
```
